# Image Memorability in the Eye of the Beholder: Tracking the Decay of Visual Scene Representations

**DOI:** 10.1101/141044

**Authors:** Melissa Le-Hoa Võ, Zoya Bylinskii, Aude Oliva

## Abstract

Some images stick in your mind for days or even years, while others seem to vanish quickly from memory. We tested whether differences in image memorability 1) are already evident immediately after encoding, 2) produce different rates of forgetting, and 3) whether the retrieval of images with different memorability scores affords differential degrees of cognitive load, mirrored by graded pupillary and blink rate responses. We monitored participants’ eye activity while they viewed a sequence of >1200 images in a repetition detection paradigm. 240 target images from 3 non-overlapping memorability classes were repeated at 4 different lags. Overall, performance decreased log-linearly with time. Differences in memorability were already visible at the shortest lag (~ 20 sec) and became more pronounced as time passed on. The rate of forgetting was steepest for low memorable images. We found that pupils dilated significantly more to correctly identified targets than correctly rejected distractors. Importantly, this “pupil old/new effect” increased with increasing number of lags and decreasing image memorability. A similar modulation of blink rates corroborated these findings. These results suggest that during memory retrieval of scenes, image inherent characteristics pose differential degrees of cognitive load on observers as seen in their eyes.

---

Chances are that the image of the Beatles crossing Abbey Road has burned itself into your memory and has hardly faded since you first saw it. This experience is backed by previous studies showing that memory for objects and scenes is massive (Brady and Oliva, 2008; Hollingworth, 2006a; Konkle et al., 2010a,b) and that image details can be stored for hours or even days (Brady and Oliva, 2008; Hollingworth, 2006a,b; Konkle et al., 2010a,b). But not all memories are created equal: while some images stick in your mind, others are forgotten immediately.

Isola et al. (2014) recently characterized the memorability of an image as the probability that an observer will detect a repetition of the image amidst a stream of distractor images and provided first evidence that memorability is an intrinsic, stable property of an image that is robust across observers. In the study presented here, we used a subset of Isola’s database of images to test whether differences in memorability are evident right after an image’s initial exposure and whether the rate of decay is a function of an image’s intrinsic memorability. In addition, we monitored viewers’ pupillary responses and blink rates. Both these easily accessible, physiological measures that are not under direct control of the observer have shown to highly correlate with cognitive load and might therefore provide a useful tool to track image memorability without the need for overt responses. Before we describe the study, we briefly review the main contributions of pupil and eye blink recordings for the study of cognitive processes.

While the overt pupillary response has be linked, since the early 60s, to covert cognitive constituents like information processing and mental load (Bradshaw, 1967; Hess and Polt, 1960, 1964; Kahneman and Beatty, 1966; Janisse, 1977; Granholm and Steinhauer, 2004), recent studies have started using pupil dilations to investigate mnemonic processes (Goldinger and Papesh, 2012; Kafkas and Montaldi, 2011, 2012; Naber et al., 2013; Otero et al., 2011; Porter et al., 2007; Võ et al., 2008). For instance, the now robust finding that pupils dilate more to “old”, studied items than to “new” ones — termed the Pupil Old/New Effect (Võ et al., 2008) or PONE — seems particularly suitable to investigate the cognitive processes involved in recogniton memory. While Võ and colleagues suggested it is the cognitive effort associated with recollective processes that drives the PONE, Otero and colleagues reported a greater PONE for items encoded under deep compared to shallow orienting instructions, suggesting the PONE may reflect the strength of the underlying memory trace. To paint a clearer picture, we recorded an additional physiological measure to study recognition memory: the blink rate.

Usually, studies interested in pupil measurements disregard blinks — i.e. the closing of the eyelid — as missing data or interpolate it. Yet an independent literature has shown that blink rates are everything but random. Thus, endogenous eye blinks — in addition to pupil dilations — provide mutually exclusive, but complementary indices of information processing (Siegle et al., 2008). In particular, blink rates have shown to decrease under conditions of high visual and/or cognitive load, since visual input is disrupted during blinks and a reduced blink rate therefore supports a continuous input of visual information especially when cognitive demands are high (Neumann and Lipp, 2002).

With regard to image memorability, these findings suggest that if the correct retrieval of scene representations generated from less memorable images requires more cognitive effort, this should elicit both an increase in pupil dilations as well as a decrease in blink rate. Similarly, the increased effort to correctly retrieve scene representations generated a longer time ago should also be mirrored in pupillary and blink responses.

## Methods

### Stimulus material and experimental design

Images were taken from the Isola et al. database (Isola et al., 2014), which includes memorability scores for 2222 scenes. From these we selected 80 images each with the highest, lowest, and medium memorability scores over time, creating three non-overlapping memorability classes (examples are shown in Figure 1).

**Figure 1.**
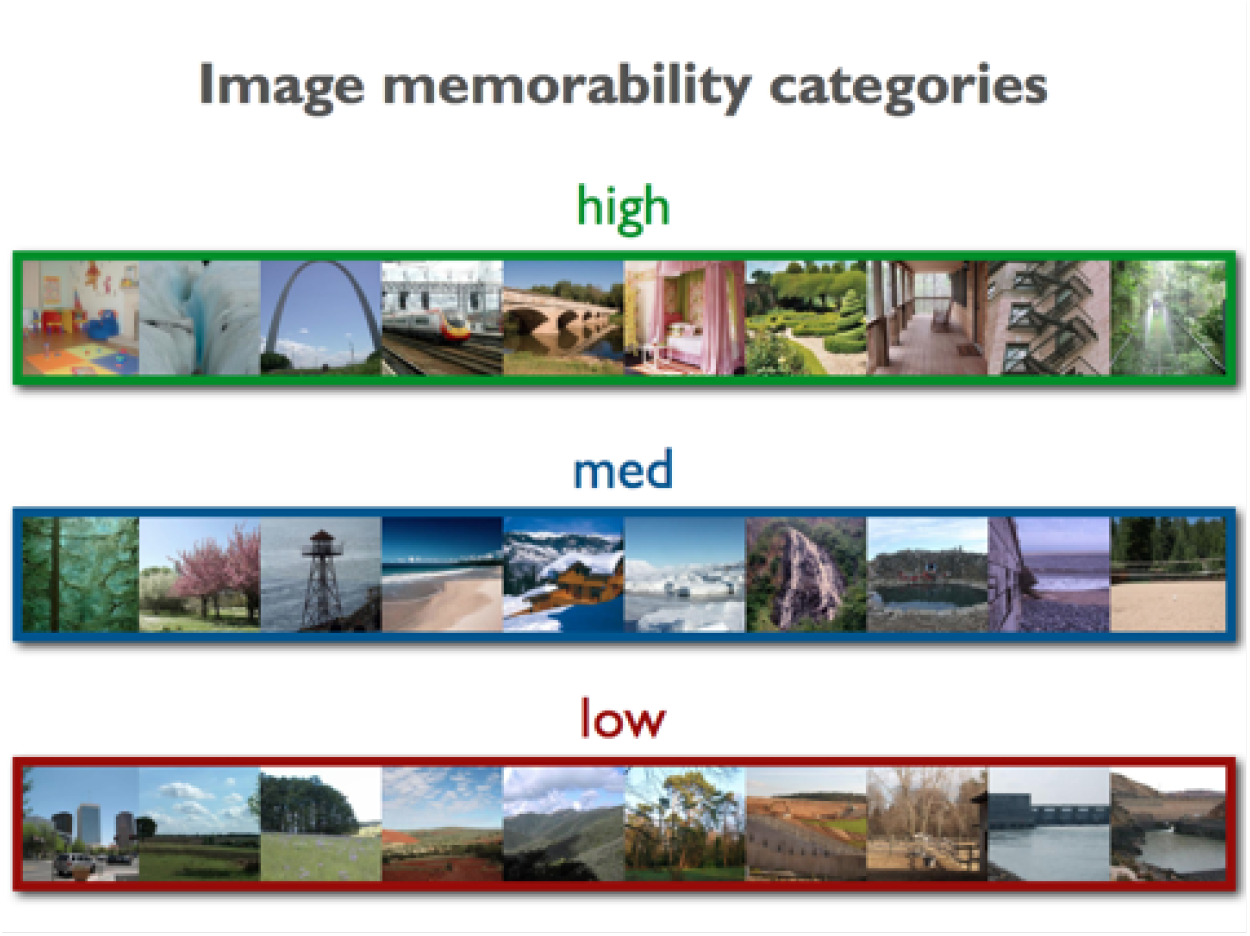
Example images from the Isola et al. database (Isola et al., 2014). We created three non-overlapping memorability classes containing 80 images each with established indices of highest, lowest, and medium memorability scores over time.

There was no difference in mean luminance between low and high memorability categories for neither LAB, HSV, nor RGB values, all *ts*(158) < 1. The experimental design included two main manipulations: image memorability (low, medium, high) and lag (8, 16, 64, 256; see Figure 2). Each participant was presented with a sequence of 1216 images, 240 of which were target images presented twice. The other 736 images were fillers, sampled randomly from the remaining images in the database. We randomly assigned target images to prespecified positions within the image sequence repeating at one of 4 lags categories (8, 16, 64, 256).

**Figure 2.**
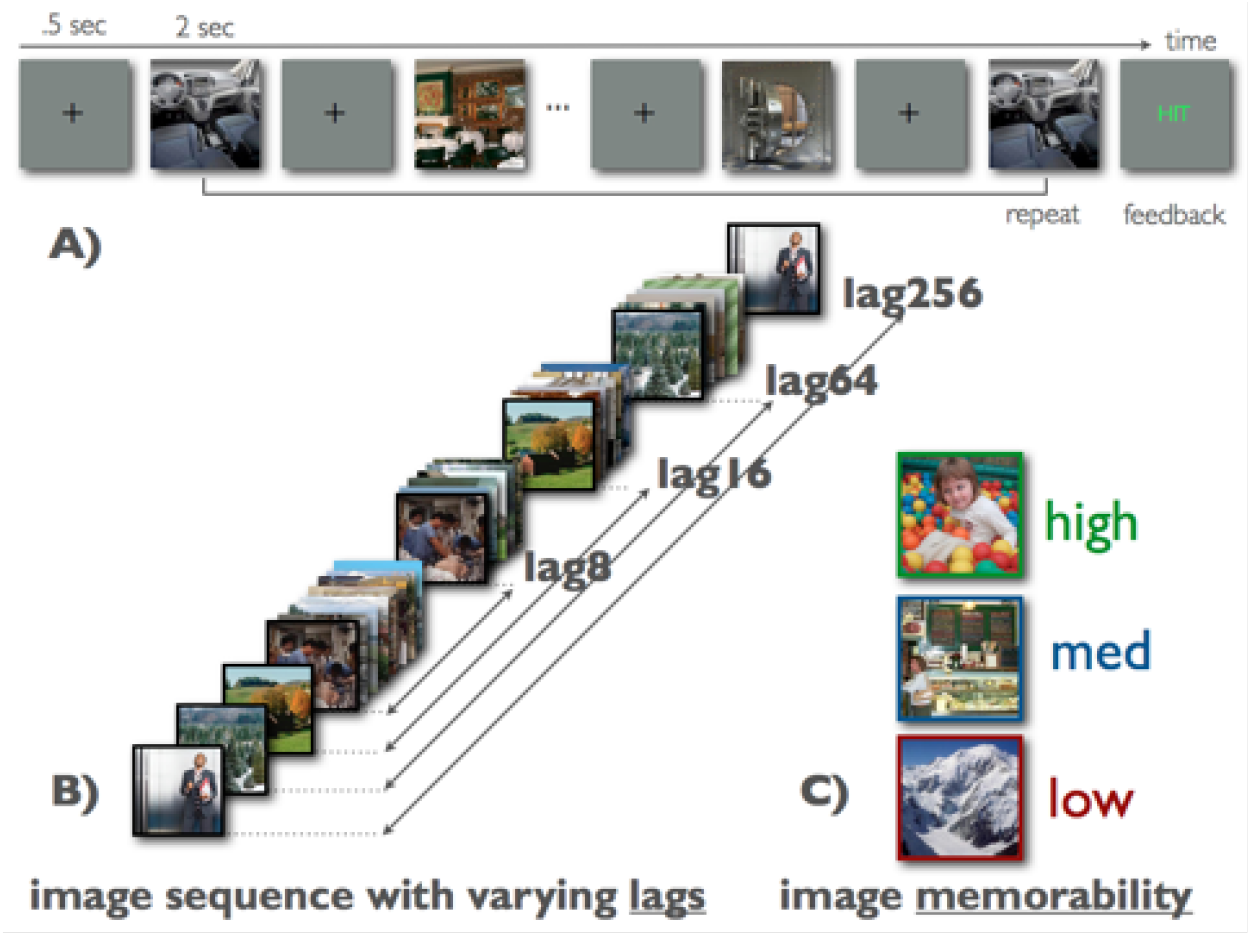
A) The sequence of image presentations used throughout the experiment consisting of 500 ms fixation cross followed by 2 sec of an image. In a repetition detection task, participants pressed a button when they detected a repeat and received feedback. B) Repetitions of target images could occur at 4 different lags (8, 16, 64, 256). C) Additionally the memorability of images varied between low, medium, and high.

#### Apparatus

Pupil dilations and blink rates were recorded with an EyeLink1000 desktop mount system (SR Research, Canada) at a sampling rate of 250 Hz. Stimulus presentation and recording of the subjects’ responses were controlled using Matlab and Psychophysics Toolbox (Brainard, 1997; Pelli, 1997). Images were presented at a distance of 55 cm on a 19-inch CRT monitor (1280x1024 pixels, 85Hz). The 512x512 pixel images subtended about 16 degrees of visual angle.

#### Participants

15 subjects (6 female) participated, ranging in age between 19 and 63 years (M=29.72, S=10.07). One subject was excluded due to faulty pupillary recordings. All were paid volunteers who gave informed consent.

#### Procedure

Participants viewed images with their head stabilized in a headrest. The experiment started with a randomized 9-point calibration and validation procedure. At regular intervals, a drift check was performed and — if necessary —recalibration took place. Before each drift check, the participant could choose to take a break. The experiment lasted 75-90 minutes.

#### Eye tracking analysis

Raw data were preprocessed using EyeLink Data Viewer (SR Research, Canada) and further analyzed using R (http://www.R-project.org/).

#### Pupil size measurement

Stimulus-locked recording segments of 2000 ms (image onset to image offset) were baseline corrected by subtracting the average dilation over a 100 ms window preceding stimulus onset, to gain comparable pupillary response indices. Peak horizontal dilations in a 2000 ms window after stimulus onset were determined and submitted separately to ANOVAs with memorability (low, medium, high) and lag (8, 16, 64, 256) as within-subject factors.

#### Blink rate

Blinks were registered by the EyeLink 1000 parser as events where the pupil size was very small, the pupil in the camera image was missing and/or severely distorted by eyelid occlusion. We determined the blink rate as the average percentage of blinks that occurred in each of the conditions.

## Results

### Performance

#### Hit Rate

A Hit was registered when the participant correctly identified the repetition of an image shown previously within the sequence. As shown in Figure 3, we found a main effect of memorability, *F*(13,2) = 63.33, *p* < .01, *pη*^2^ = .70, in that Hit rate decreased with lower image memorability. The main effect of lag, *F*(13,3) = 97.76, *p* < .01, *pη*^2^ = .63, was characterized by a decrease of performance with increasing lags. In addition we found an interaction of memorability and lag, *F*(13,6) = 10.22, *p* < .05, *pη*^2^ = .18, with differences in slopes across lags as a function of memorability. The rate of forgetting increased from high-mem images (-3.58), over medium-mem images (-6.94) to low-mem images (-8.30). Further, Hit rates already differed significantly at the shortest lag: high (97%) vs. low (71%), *t*(12) = 3.79, *p* < .05.

**Figure 3.**
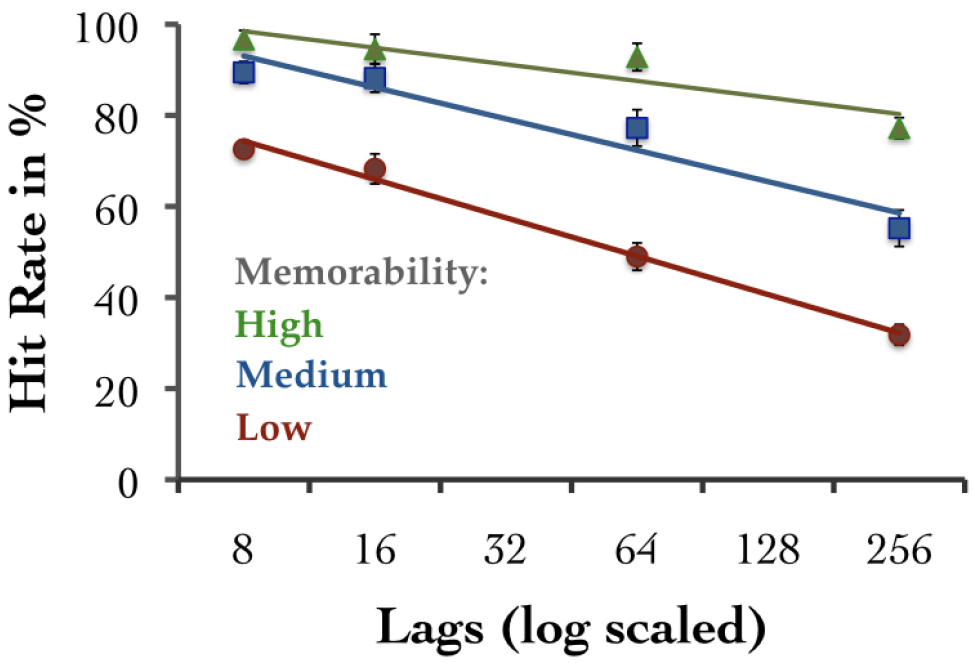
Hit rates in percent as a function of lags (log scaled: 8, 16, 64, and 256) and memorability (high = green, medium = blue, low = red). Bars depict standard errors.

#### Response Times

RTs were measured from scene onset to button press and are only shown for Hits. One subject did not score a single Hit for low memorability images at lag-256; we therefore excluded the subject for this particular analysis. Similar to Hit rates, Figure 4 shows a main effect of memorability, *F*(12,2) = 105.72, *p* < .01, *pη*^2^ = .59, amain effect of lag, *F*(12,3) = 53.18, *p* < .01, *pη*^2^ = .61, as well as an interaction of memorability and lag, *F*(12,6) = 2.39, *p* < .05, *pη*^2^ = .09, with differences in slopes across lags as a function of memorability: high = 24.07, medium = 40.08, and low = 41.91. RTs to images of varying memorability already differed significantly at the shortest lag: high (779ms) vs. low (897ms), *t*(12) = 9.48, *p* < .01.

**Figure 4.**
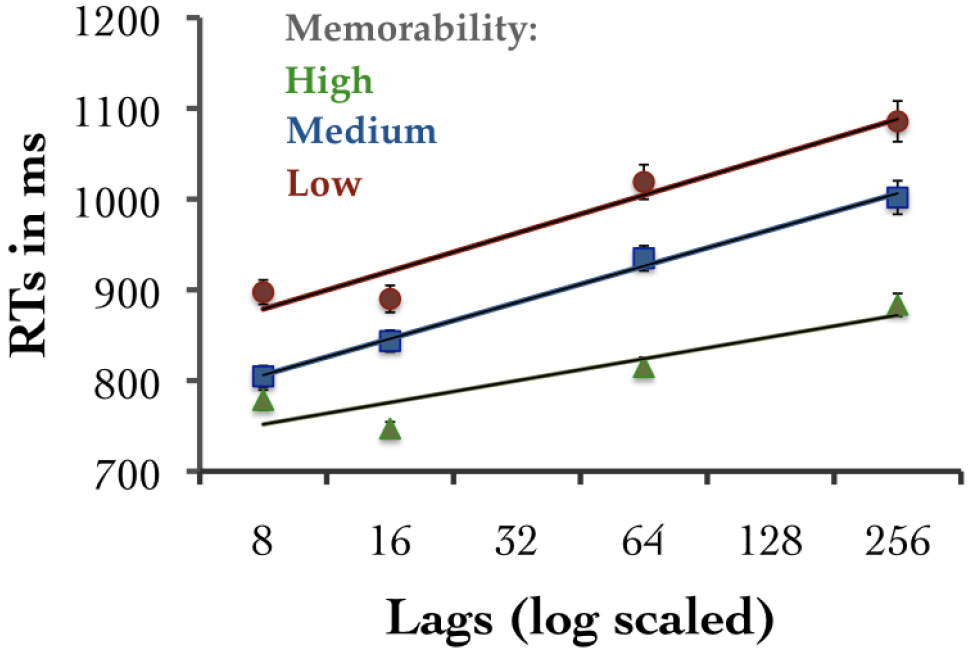
Reaction times in percent as a function of lags (log scaled: 8, 16, 64, and 256) and memorability (high = green, medium = blue, low = red). Bars depict standard errors.

### Pupil Old/New Effect

The Pupil Old/New Effect (PONE) is measured as the difference in pupillary responses (i.e. peak horizontal dilations) to old images correctly identified as “old” (Hits) versus new images correctly identified as “new” (Correct Rejections, CRs).

#### Image Memorability

As can be seen in Figure 5, there was a significant PONE during retrieval in that pupils dilated more to Hits vs. CRs (*M* = 241, *t*(13) = 5.98, *p* < .01). In addition, differences in image memorability were mirrored in a graded PONE response, *F*(13,2) = 3.33, *p* < .05, *pη*^2^ = .20, with low-mem images eliciting a greater PONE than high-mem images, *t*(13) = 2.23, *p* < .05.

**Figure 5.**
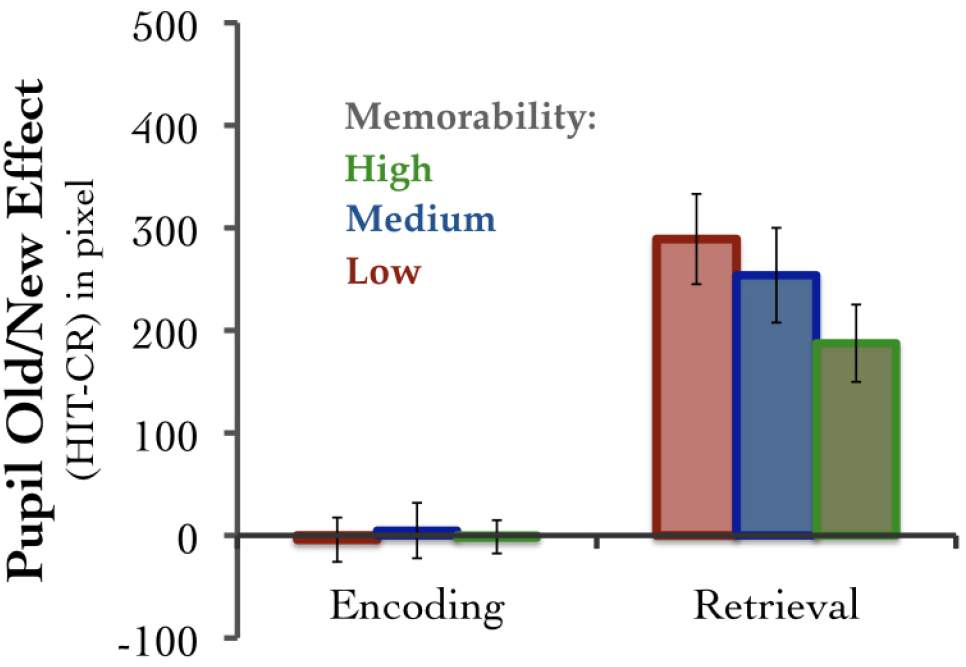
The Pupil Old/New Effect — measured as the difference in pupillary responses (i.e. peak horizontal dilations) to Hits versus Correct Rejections (CRs) — as a function of image memorability (high = green, medium = blue, low = red) both during encoding (left) and retrieval (right). Bars depict standard errors.

#### Encoding-Retrieval Lag

The PONE was also modulated by the lag between image encoding and its successful retrieval, *F*(13,3) = 3.98, *p* < .05, *pη*^2^ = .23, characterized by an increased PONE at the longest lag compared to lag-16, *t*(13) = 2.57, *p* < .05 (see Table 1).

**Table 1.** The Pupil Old/New Effect [SE] in pixel as a function of lag (8, 16, 64, 256).

| lag | PONE Mean | PONE SE |
| --- | --- | --- |
| 8 | 224 | 28 |
| 16 | 185 | 39 |
| 64 | 210 | 37 |
| 256 | 304 | 59 |

### Blink Old/New Effect

The Blink Old/New Effect (BONE) was calculated as the difference in mean blinking rate during the viewing of old images correctly identified as “old” (Hits) versus new images correctly identified as “new” (CRs).

#### Image Memorability

As shown in Figure 6, there was a significant BONE characterized by reduced blinking rates for Hits vs. CRs during retrieval (*M* = −16%, *t*(13) = 3.47, *p* < .01). The BONE during retrieval was further modulated by image memorability, *F*(13,2) = 6.04, *p* < .01, *pη*^2^ = .32, with significantly lower blink rates for low-mem compared to high-mem images, *t*(13) = 2.83, *p* < .05.

**Figure 6.**
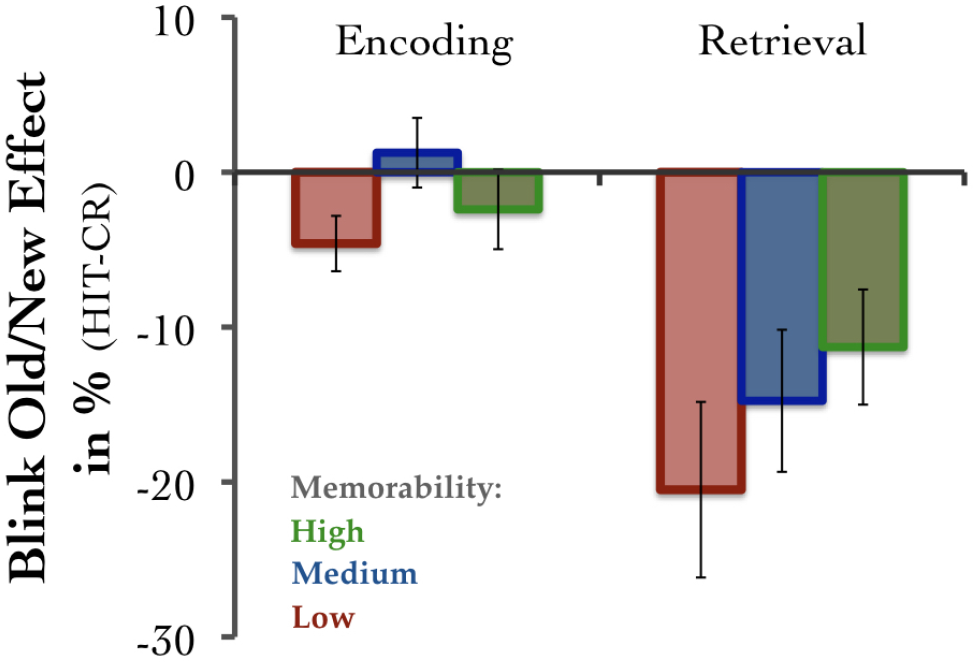
The Blink Old/New Effect in % — measured as the difference in blink rates to Hits versus Correct Rejections (CRs) — as a function of image memorability (high = green, medium = blue, low = red) both during encoding (left) and retrieval (right). Bars depict standard errors.

#### Encoding-Retrieval Lag

The BONE was not significantly modulated by the lag between image encoding and its successful retrieval, *F*(13,3) = 1.98, *p* = .13, *pη*^2^ = .13, but there was a marginally significant decrease of blink rate at lag-256 compared to lag-16, *t*(13) = 2.13, *p* = .05.

Note that the very same images did not show either a PONE or a BONE during their encoding (*M* = −.24, *t* < 1 and *M* = −2%, *t* = 1.11, *p* = .29, respectively), thus the old/new effects and their modulations by memorability observed during retrieval are not due to mere differences in visual features, but reflect actual memory retrieval processes.

## Discussion

Image memorability has strong and robust effects on both recognition memory performance and eye activity measures. This study not only corroborated earlier findings according to which memorability is an intrinsic property of an image that is shared across different viewers and remains stable over time (Isola et al., 2014), but also clearly showed that low memorable images compared to highly memorable images 1) show a decline in recognition performance of over 25% only 20 seconds after initial presentation, 2) produce steeper rates of forgetting, and 3) are accompanied by increased pupillary responses and decreased blink rates.

While it is not surprising that some images are less memorable than others, it does seem striking how quickly differences in image memorability become evident. In addition to the almost instantaneous decline in memory performance, mental representations of less memorable images also faded away at a faster rate. After only 10 minutes, recognition memory for low-memorable images dropped dramatically from 71% to only 32%, while recognition memory for highly memorable images merely declined from 97% to 78% in the same time interval. Moreover, the successful retrieval of scene representations from low memorability images involved more more cognitive effort, as seen in prolonged RTs, greater pupil dilations, and decreased blink rates. However, we found no modulation of pupillary responses during encoding, neither by memorability nor by subsequent successful retrieval (but see Naber et al. (2013) for evidence that pupil size can predict success or failure of memory formation during encoding).

The pupillary response has been widely studied in the cognitive domain and recently gained momentum in the investigation of recognition memory (Goldinger and Papesh, 2012; Kafkas and Montaldi, 2011, 2012; Naber et al., 2013; Otero et al., 2011; Porter et al., 2007; Võ et al., 2008). While it is agreed upon that pupils generally dilate more under high cognitive load, it is not quite clear what cognitive processes are reflected in the pupil response during memory retrieval. While Võ and colleagues suggested that the Pupil Old/new Effect (PONE) mirrors the cognitive effort associated with recollective processes (Võ et al., 2008), Otero and colleagues have argued that the PONE reflects the strength of the underlying memory trace (Otero et al., 2011). These two views make opposing, but testable claims with regard to the memorability of images: If greater memory strength per se would elicit greater pupil dilations, highly memorable images should be accompanied by a greater PONE. If the PONE, on the other hand, reflects the cognitive effort involved in retrieving memory representations, highly memorable images should elicit the smallest pupillary response. The data reported here provide evidence for the latter view.

This was further backed by the finding that blink rates — which have shown to decrease under higher cognitive load (Neumann and Lipp, 2002) — were lowest for low as compared to high memorable images, signaling increased cognitive load required to successfully retrieve less memorable images. Overall, blink rates decreased during the correct recognition of “old” images as compared to “new” images. We therefore propose that this “Blink Old/New Effect” (BONE) could be used as a complementary measure for the investigation of memory processes. Siegle and colleagues have previously reported that pupil dilations reflect sustained information processing, whereas blinks occur during early sensory processing and following sustained information processing (Siegle et al., 2008). Importantly, their data support our findings showing that during stimulus presentation a higher cognitive load leads to an increase in pupillary response and a simultaneous decrease in blink rate (see Siegle et al. (2008), Figures 1a,b).

In sum, we have shown that the intrinsic memorability of an image has both immediate and long-lasting effects on recognition performance and can be tracked using two easily accessible and complementary physiological measures: the pupillary response and endogenous blink rate. Image memorability is therefore indeed mirrored in the eye of the beholder.

## Acknowledgements

This work was funded by a research award from Google and a National Science Foundation grant (1532591) to A.O, as well grant a grant from the Deutsche Forschungsgemeinschaft (DFG) VO 1683/2-1 to M.L.V.

